# Identification of functional long non-coding RNAs in *C. elegans*

**DOI:** 10.1101/383919

**Authors:** Alper Akay, David Jordan, Isabela C. Navarro, Tomasz Wrzesinski, Chris P. Ponting, Eric A. Miska, Wilfried Haerty

## Abstract

**Background:** Functional characterisation of the compact genome of the model organism *Caenorhabditis elegans* remains incomplete despite its sequencing twenty years ago. The last decade of research has seen a tremendous increase in the number of non-coding RNAs identified in various organisms. While we have mechanistic understandings of small non-coding RNA pathways, long non-coding RNAs represent a diverse class of active transcripts whose function remains less well characterised.

**Results:** By analysing hundreds of published transcriptome datasets, we annotated 3,397 potential lncRNAs including 146 multi-exonic loci that showed increased nucleotide conservation and GC content relative to other non-coding regions. Using CRISPR / Cas9 genome editing we generated deletion mutants for ten long non-coding RNA loci. Using automated microscopy for in-depth phenotyping, we show that six of the long non-coding RNA loci are required for normal development and fertility. Using RNA interference mediated gene knock-down, we provide evidence that for two of the long non-coding RNA loci, the observed phenotypes are dependent on the corresponding RNA transcripts.

**Conclusions:** Our results highlight that a large section of the non-coding regions of the *C. elegans* genome remain unexplored. Based on our in vivo analysis of a selection of high-confidence lncRNA loci, we expect that a significant proportion of these high-confidence regions is likely to have biological function at either the genomic or the transcript level.

## Background

Transcription is not limited to the protein-coding regions of eukaryotic genomes, but instead has been observed to be pervasive in all organisms that have been studied so far. As a consequence of transcriptional activity over non-coding sections of the genomes, tens of thousands of short, <200 nucleotide (nt), and long (>200 nt) non-coding RNAs have now been annotated [1,2]. While much is known about the biological role of most classes of small non-coding RNAs (e.g microRNA, Piwi-associated RNA, small nucleolar RNA, small interfering RNA) [3–5], relatively little is known about long non-coding RNAs (lncRNAs). Whether most eukaryotic lncRNAs are functional has long been debated because of their low expression levels and rapid evolutionary turnover when compared to protein coding genes [6,7]. However, the molecular activities of more than a hundred of such loci have now been described [8–12] including many that appear to regulate the expression of protein-coding genes. Only a small proportion of these loci have been demonstrated to be fundamental to eukaryote biology from mutations that affect their expression or function leading to severe developmental defects or to lethal phenotypes (for example,[13,14]). While transcription of some lncRNAs has been shown to originate at promoter or enhancer elements with potential DNA-dependent function [15], the activity of others depends on the RNA transcript, acting either in *cis* or *trans; e.g* targeting protein complexes to chromatin or directly interacting with other RNAs, including mRNAs, lncRNAs, or microRNAs [16]. The proportions of lncRNAs belonging to each functional class remain unknown owing to painstaking experimental validations, including both knockout and knockdown assays being required.

*C. elegans* has been invaluable for the discovery of multiple non-coding RNA pathways and is an important model organism for genetic studies. Nevertheless, only one study has yet sought to identify and annotate lncRNAs in *C. elegans,* resulting in 801 annotated loci, of which only 170 had evidence of polyadenylation [17]. Furthermore, experimental characterisation of *C. elegans* lncRNAs has been limited [18–20]. Despite evidence confirming their expression [20], previously identified lncRNAs of *C. elegans* still lack functional validation [17].

Using publicly available RNA-Seq libraries representing diverse *C. elegans* developmental stages, we sought to annotate novel expressed long non-coding loci and to characterize informative features such as nucleotide composition, evolutionary conservation, transcript expression and functional enrichment. To assess the physiological impact of mutations within these lncRNAs and thus the biological importance of these loci, we used CRISPR/Cas9 to generate large genomic deletions for ten lncRNA loci. Six of these intergenic lncRNA loci yielded significant phenotypes upon deletion, and at least two of these have physiological functions that are RNA-dependent. Our study and associated experimental validation demonstrate that physiological lncRNA function in nematodes can be RNA- and/or transcription-dependent. Furthermore, we extrapolate that a significant proportion of the newly identified multi-exonic non-coding loci in the *C. elegans* genome might be functional at the genomic or the transcript level.

## Results

### Long non-coding RNA annotation in *C. elegans*

We investigated 209 publicly available RNA-Seq datasets from diverse developmental stages (Additional File 1) to annotate de novo non-coding transcripts in *C. elegans.* After filtering for size, coding potential, overlap with existing protein coding genes and loci found in close proximity in the same orientation to annotated genes (see Methods), we identified 3,397 long (>200 nt) non-coding RNAs expressed across *C. elegans* development (Additional file 2). Only 197 of these loci were annotated previously [17].

In total, 146 multi-exonic and 3,251 mono-exonic loci were identified (18 and 179 of which were previously known [17]). CAGE data [21] was then used to accurately annotate transcriptional start sites, and ChlP-Seq [22] and CLIP-Seq [23] data were used to identify transcription factor and AGO binding sites within the loci (Additional file 3). As observed in all other model organisms, the identified lncRNA loci are smaller than annotated protein coding genes and are expressed at significantly lower levels (Additional file 4). LncRNA exons also tend to have a GC content that is lower than protein coding sequences but higher than intronic sequences (Fig. 1a, b) as observed previously for other eukaryotes[24]. Inter-species sequence conservation for multi-exonic lncRNAs was lower than for protein-coding genes but higher than for mono-exonic lncRNAs (Kruskal Wallis test, P=5.73×10”^5^, Fig. 1c). Our newly annotated multi-exonic lncRNAs show sequence features similar to the final set of 170 lncRNA reported by Nam and Bartel [17], (Kruskal Wallis test, P=0.79 and P=0.15 for nucleotide conservation and composition respectively) showing the complementarity of these lncRNA annotations.

**Fig.1.**
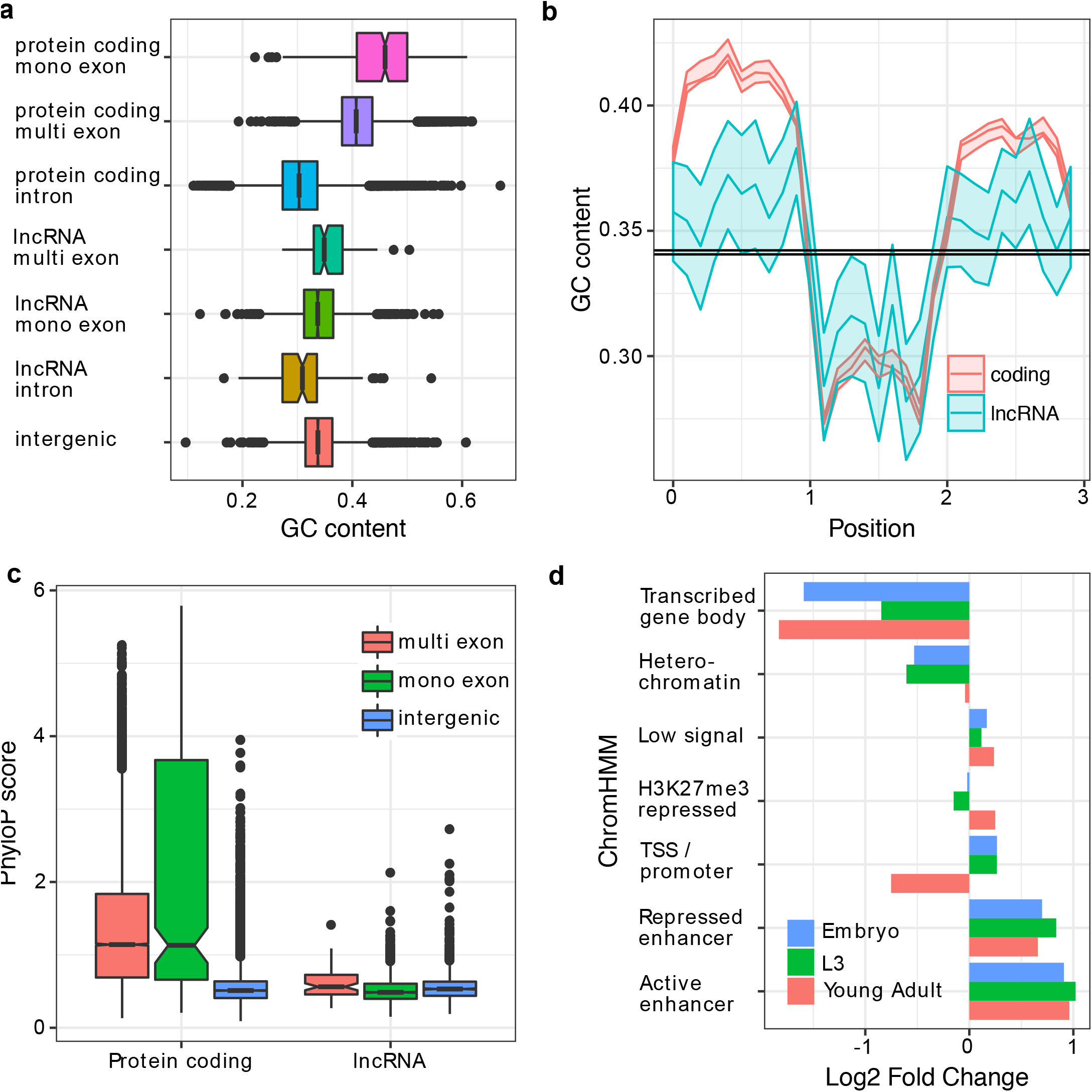
LncRNAs sequence features in C. elegans **a.** Nucleotide compositions of exons and introns of lncRNAs and protein coding genes classified according to their gene model. **b.** GC content variation within non overlapping windows each representing 10% of the sequences across multiexonic proteincoding and lncRNA loci, the black band represent the GC content of flanking intergenic sequences. **c.** Nucleotide conservation (Phylo score) comparison among intergenic sequences, multior monoexonic lncRNAs and protein coding loci. **d.** Enrichment of lncRNAs for chromatin annotations identified by Daugherty et al [27]. Transcribedgene body: ensemble of ChromHMM states characterized by H3K79me2, H3K36me3, H3K4me1, and H4K20me1. Repressed enhancers: ensemble of ChromHMM states characterized by H3K4me1 and H3K27me3. Low signal: regions without histone modification signals.

Distinct chromatin states inferred from histone modifications using ChromHMM [25] have been shown to associate with specific genomic elements (for example, transcriptional start sites and promoters, transcriptional elongation and gene bodies, enhancers, transposable element-derived sequences). Using previously published chromatin annotations in *C. elegans* [26,27], we assessed the functional enrichment of our newly annotated lncRNA at each of these annotated genomic elements. Enhancers, identified either by Evans et al. [26] (1.7 fold enrichment P<0.0001) or Daugherty et al. [27] (2.0 fold enrichment, P<0.0001), significantly overlapped with these lncRNA loci but chromatin states associated with transcription elongation (“transcribed gene body”) were depleted at all developmental stages (Figure 1d, Additional File 5). These results could be explained by the observed low expression level of the lncRNAs. Our results are in agreement with Evans et al. [26], who showed that the chromatin states reflecting transcription elongation were associated with the most highly expressed genes in their study. Active enhancers were particularly enriched within single exon lncRNAs at all developmental stages (2.0, 2.6 and 2.4 fold enrichment respectively at the early embryonic, L3, and young adult stages, P<0.001 in all comparisons, Additional File 5). This result is consistent with enhancer RNAs rarely being spliced [28]. In contrast, multi-exonic loci were only enriched for active enhancers during early embryonic stage (2.3 fold enrichment, P<0.0001) which likely reflects the fewer number of multi-exonic lncRNA expressed at later stages.

Half of all lncRNA loci are expressed in at least 12 libraries (FPKM > 1; or at least 41 libraries if FPKM>0.1; Fig. 2a). This restricted expression could reflect that many of the newly annotated loci are the result of transcriptional noise, and therefore likely non-functional [15]. However, many of the remaining lncRNAs, most specifically multi-exonic lncRNAs, appear to be expressed in a tissue- and stage-specific manner (Fig. 2b). 26 lncRNAs (10 multi-exonic) were expressed in more than 90% of the libraries (≥188 libraries). Highly reproducible loci (≥100 libraries) tended to have a significantly higher sequence conservation (Kruskal Wallis test, corrected P=0.0077) and higher GC content (Kruskal Wallis test, P=3.5×10”^4^ after Bonferroni correction) compared with loci with limited reproducibility (<10% libraries) (Fig. 2c, 2d). Highly reproducibly expressed loci also tended to have stronger enrichment for enhancer regions identified in embryos (2.2-3.1 fold enrichment) or in L3 larvae (2.7-4.4 fold enrichment, Additional File 5).

**Fig.2.**
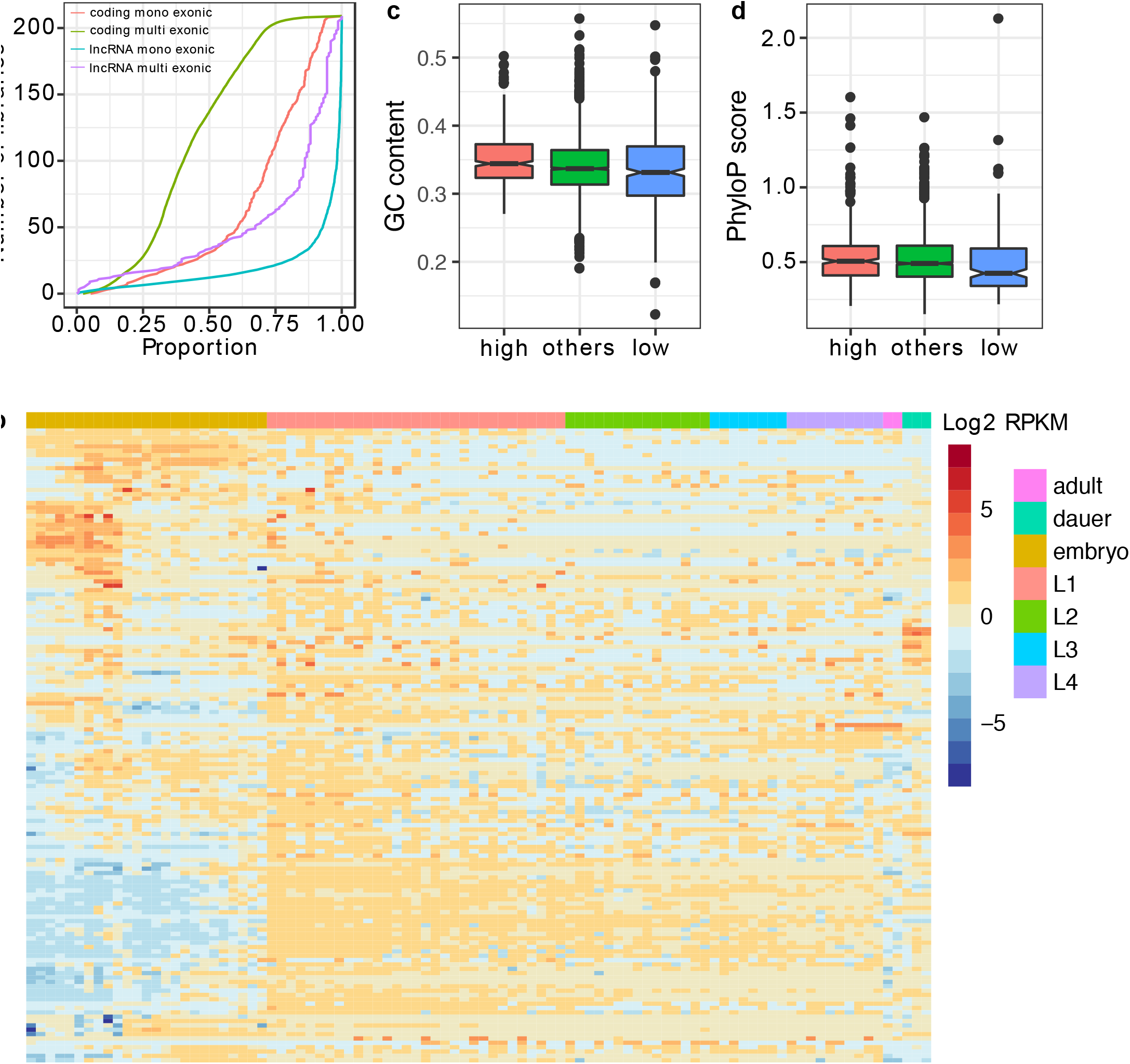
LncRNA expression properties. **a.** Cumulative distribution of the proportion of multi-exonic, mono-exonic lncRNA and protein coding loci identified as expressed across all libraries. **b.** Expression (log2 RPKM) across *C. elegans* development of 146 multi-exonic lncRNAs. Each column represents the average expression at one time point for whole individuals in standard conditions. GC composition (**c**) and nucleotide conservation (**d**) for lncRNA loci depending on the reproducibility of lncRNA model predictions across libraries. Low: ≤ 20 of the libraries (low), others: 20 to 100 (others), high: ≥ 100 libraries.

### Functional characterisation of lncRNA loci

Higher sequence conservation of the 146 multi-exonic lncRNA loci, together with their higher exonic GC content and their splicing, could reflect organismal function. To test this hypothesis, we used CRISPR/Cas9 genome editing to generate targeted deletions in 10 of the multi-exonic lncRNA loci that each showed high sequence conservation, high expression, evidence from multiple libraries and that are not overlapping with neighbouring coding regions. We were successful in generating large genomic deletions for ten of these lncRNA loci (Table 1) out of twenty that were initially targeted. This success rate was due mostly to the limited efficiency of plasmid based protocols that were available at the time, as compared to direct protein / RNA injection methods developed later [29]. Of these lncRNA locus deletions, nine removed at least one exonic region and one removed a region just 5’ of a lncRNA locus (Additional file 6).

**Table 1.**
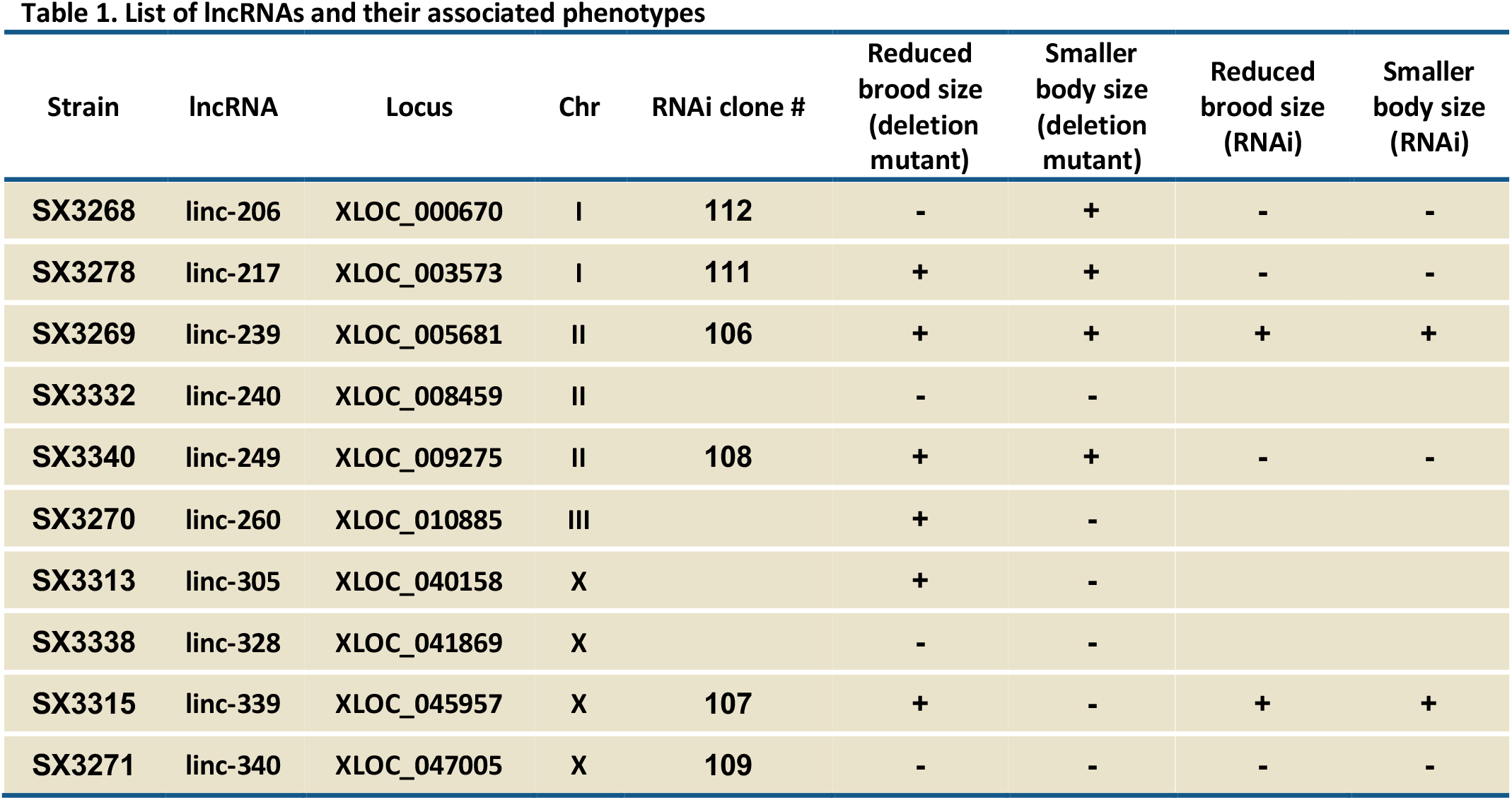
List of lncRNAs and their associated phenotypes

| Strain | lncRNA | Locus | Chr | RNAi clone # | Reduced brood size (deletion mutant) | Smaller body size (deletion mutant) | Reduced brood size (RNAi) | Smaller body size (RNAi) |
| --- | --- | --- | --- | --- | --- | --- | --- | --- |
| **SX3268** | linc-206 | XLOC_000670 | I | **112** | - | + | - | - |
| **SX3278** | linc-217 | XLOC_003573 | I | **111** | + | + | - | - |
| **SX3269** | linc-239 | XLOC_005681 | II | **106** | + | + | + | + |
| **SX3332** | linc-240 | XLOC_008459 | II |  | - | - |  |  |
| **SX3340** | linc-249 | XLOC_009275 | II | **108** | + | + | - | - |
| **SX3270** | linc-260 | XLOC_010885 | III |  | + | - |  |  |
| **SX3313** | linc-305 | XLOC_040158 | X |  | + | - |  |  |
| **SX3338** | linc-328 | XLOC_041869 | X |  | - | - |  |  |
| **SX3315** | linc-339 | XLOC_045957 | X | **107** | + | - | + | + |
| **SX3271** | linc-340 | XLOC_047005 | X | **109** | - | - | - | - |

All 10 lncRNA deletion mutants initially failed to display overt, gross phenotypes such as sterility, embryonic lethality or abnormal body development. To undertake a more extensive characterisation, we captured the development of the mutant animals alongside wild type control animals using an automated microscopy system. This system records the development of multiple animals simultaneously, and permits phenotypic analysis in an unbiased manner. Two phenotypes that can influence the life-history and fitness of populations [30], brood size and growth rate, were selected for the automated analysis. Six of 10 lncRNA deletion mutants (linc-217, linc-239, linc-249, linc260, linc-305 and linc-339) yielded significantly reduced brood size (Fig.3a) and 4 of 10 mutants (linc-206, linc-217, linc-239 and linc-249) displayed reduced growth rate (slower body size increase) over development (Fig.3b). Three mutants (linc-217, linc-239 and linc-249) showed alterations of both phenotypes (Table 1, Fig. 3).

**Fig.3.**
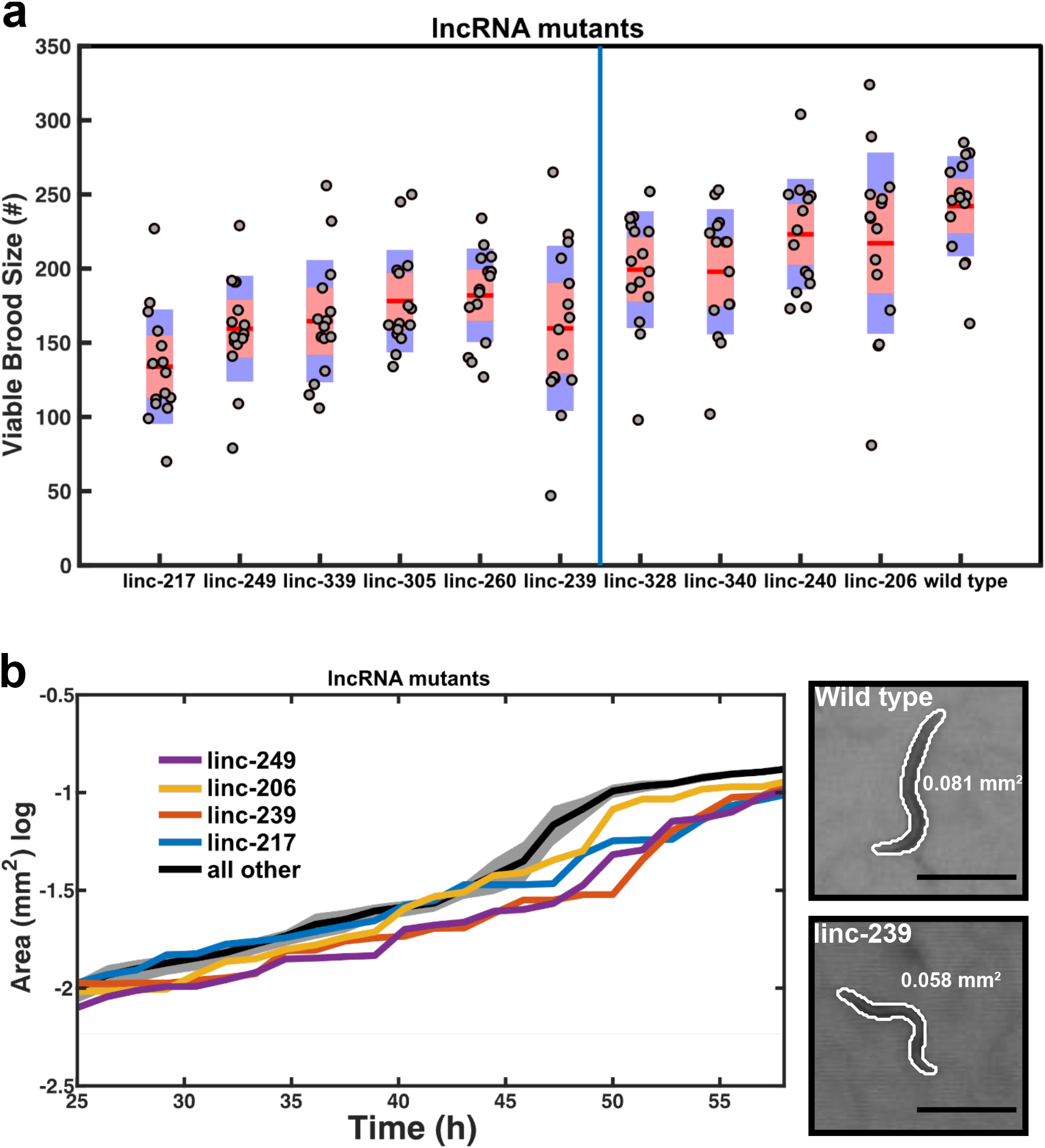
Phenotyping of 10 lncRNA deletion mutants. **a** Brood sizes are presented with their standard deviations (blue area) and the 95% confidence interval of the mean (red area). Samples were compared to wild type animals using a pairwise 2-sample t-test with a multiple test (Bonferroni) correction. Samples are ordered by increasing p-value and those found to be significant at (p≤0.05) are shown to the left of the blue line (n=15 animals / mutant). **b** Growth curves were compared to wild type animals and those found not to be significantly different are shown by their mean across strains (black line) with the standard error of the mean (grey area). Those found to be significantly different from the control are shown individually as means only. Inset shows example images of the wild type (top) and linc-239 mutant (bottom) at 45 hours post hatching with the computer-generated outlines, and computed area (black line=500μm).

These phenotypes could be due to the removal of either the lncRNA transcript or of the genomic locus which, in some instances, harboured annotated transcription factor binding and enhancer sites (Additional file 3). To distinguish between these two possibilities, we generated dsRNA expression vectors for RNAi targeting of the lncRNA transcripts in wild type animals (Table 1). Using four biological replicates per assay, we targeted 6 lncRNA transcripts using RNAi. We left out linc-240, linc-328, linc-260 and linc-305 because these were either lacking any phenotype or yielded only a weak phenotype when deleted.

Of the four of these six lncRNA loci whose deletion yielded a brood size phenotype, two (linc-239 and linc-339) yielded an equivalent phenotype when expression was reduced using RNAi (Fig. 4a); for one of these lncRNA loci, linc-239, equivalent reduced growth rate phenotypes were observed for both its knockout and knock-down (Table 1, Fig. 4b). Direct comparison of the phenotypes between lncRNA deletion mutants and RNAi knock-down of lncRNAs shows that RNAi knock-down phenotypes are slightly weaker or equivalent to deletion mutants (Fig. 5a,b). Expression of the two lncRNAs, linc-239 and linc-339, was validated by RT-PCR (Additional file 7) and we calculated their expression to be highest during larval development in comparison to embryogenesis (Fig. 5c). This indicates that, for these two loci, the phenotypes are caused by the disruption of their RNA transcript-dependent functions. RNAi targeting of another lncRNA, linc-339, also showed a reduced growth rate, a phenotype that was not observed in the deletion mutant, which could therefore be a consequence of off-target effects (Fig. 4b). Three additional lncRNA strains (linc-206, linc-217, linc-249) yielded discordant phenotypes when disrupted or subjected to RNAi (Table 1; Fig. 4a). This would be consistent with functions of these loci being RNA-independent.

**Fig.4.**
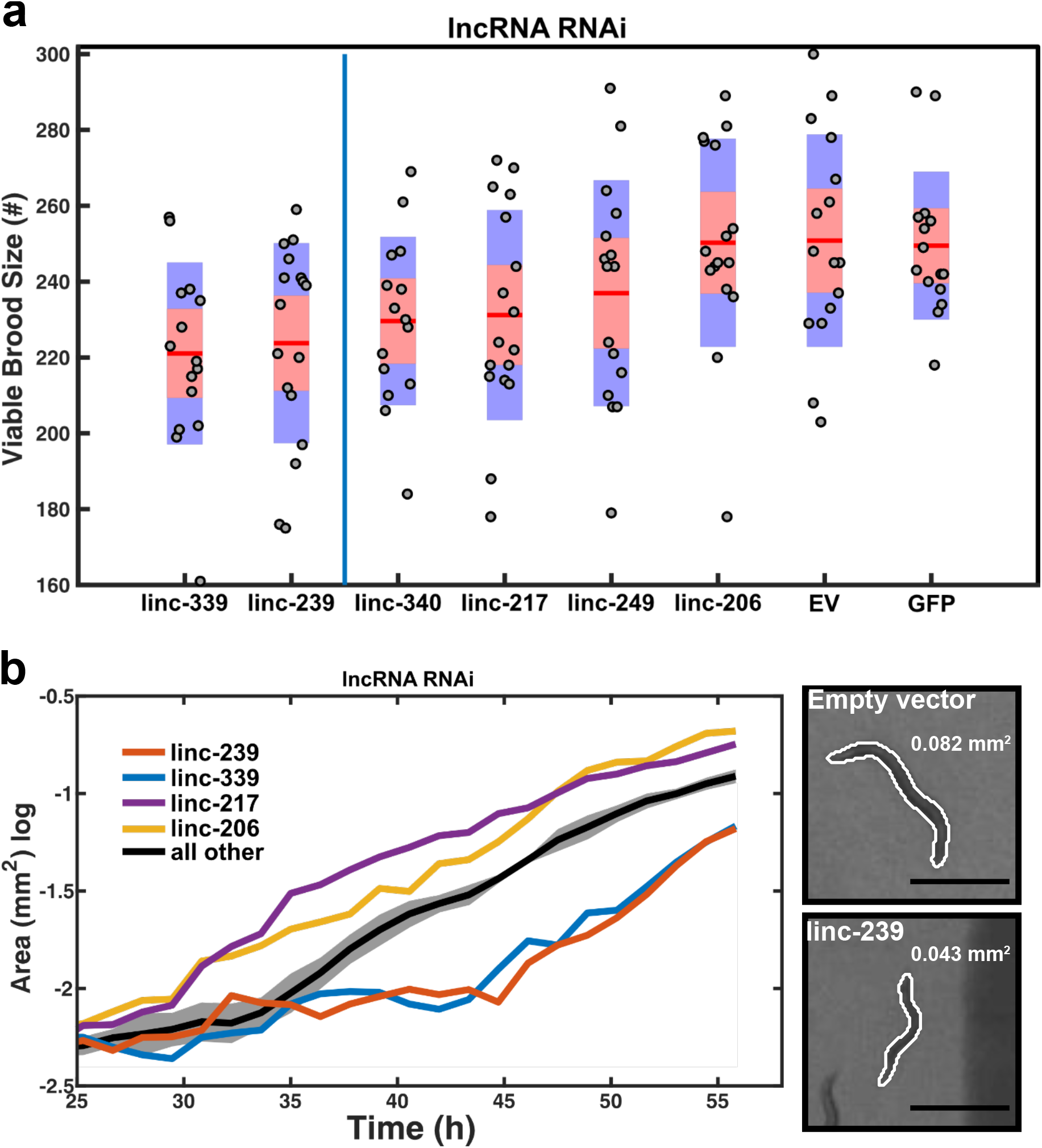
RNAi mediated knock-down of lncRNAs. **a** Brood sizes are presented with their standard deviations (blue area) and the 95% confidence interval of the mean (red area). Samples were compared to “empty vector” control animals using a pairwise 2-sample t-test with a multiple test (Bonferroni) correction. Samples are ordered by increasing p-value and those found to be significant at (p≤0.05) are shown to the left of the blue line (n=18 animals / mutant). Empty vector (EV), GFP and linc-340 RNAi are negative controls. **b** Growth curves were compared to “empty vector” animals and those found to be not significantly different are shown by their mean across strains (black line) with the standard error of the mean (grey area). Those found to be significantly different from the control are shown individually as means only. Inset shows example images of the “empty vector” (top) and linc-239 RNAi (bottom) at 45 hours post hatching with the computer-generated outlines, and computed area (black line=500μm).

**Fig 5.**
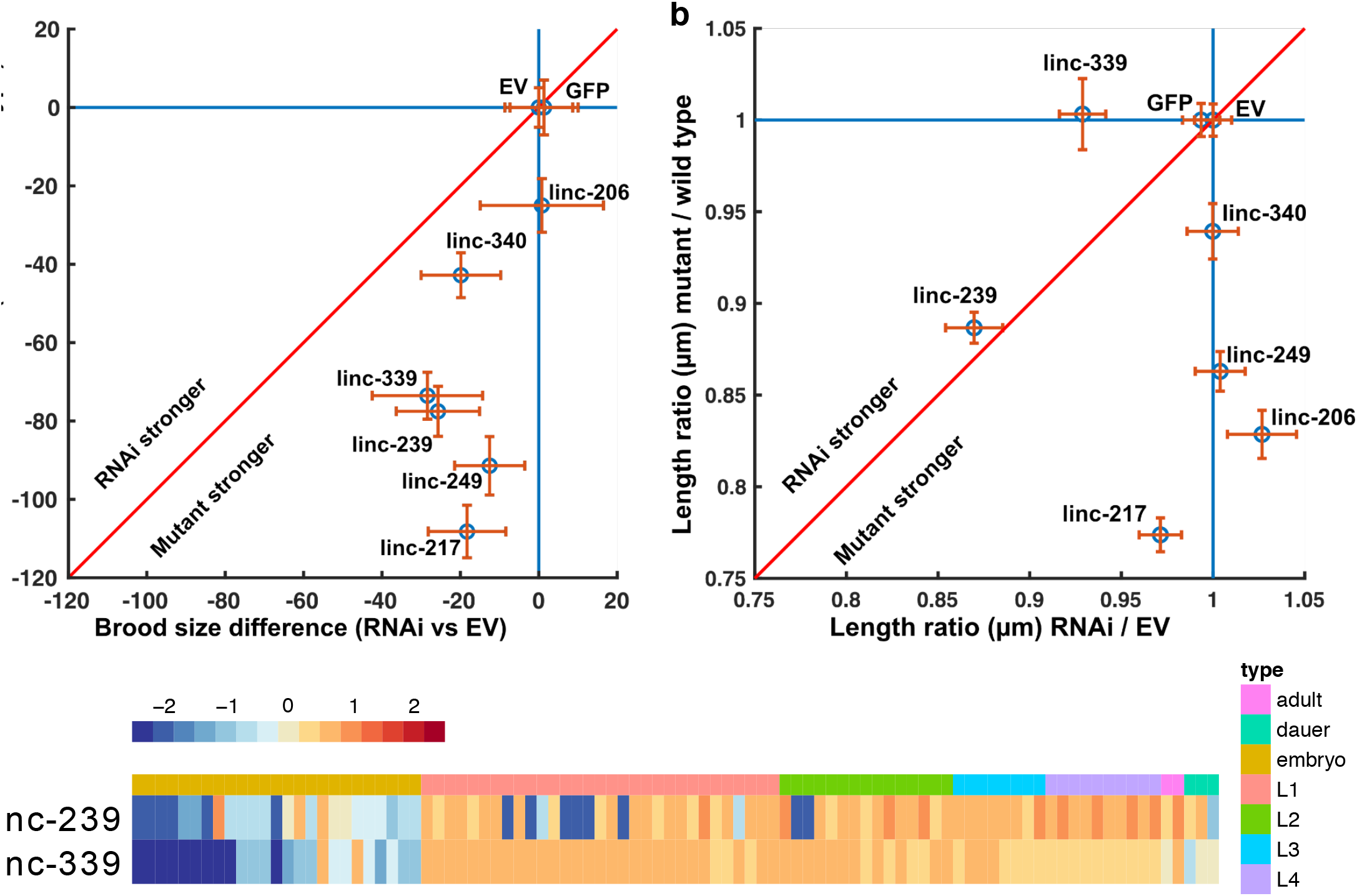
Comparison of phenotypes arising from lncRNA genomic deletion mutants and phenotypes arising from RNAi-mediated knock-down of lncRNA transcript. To compare the effects of the disruption of a lncRNA genomic locus to the knockdown of the corresponding lncRNA transcript by RNAi, the mean brood size reduction compared to the control (**a**), and the ratio of the length at 50 hours relative to the control (**b**), were plotted. These are shown as a scatter plot of the mean reduction (**a**, blue circle) or the mean ratio (**b**, blue circle) with the 95% confidence interval of the mean (orange lines). If the mutations or the RNAi yield an effect, data fall below the line y=1 and to the left of x=1. If mutants and RNAi yield similar effects, data fall along the red line; above the red line indicates that RNAi has a greater effect, while below the red line indicates that the genomic mutation has a greater effect. **c.** Expression (log2 RPKM) across *C. elegans* development for linc-239 and linc-339.

## Discussion

The identification of functional non-coding elements, including transcribed non-coding sequences, in genomes has long relied on computational predictions based on sequence conservation [31], or biochemical activity [32]. However regardless of the preferred approach to predict functional sequences, only experimental validation can truly substantiate the inferred functionality of an element.

In our study, we first provide a novel annotation of intergenic lncRNAs in *C. elegans.* This work expands on the previous annotations delivered by Nam and Bartel [17] as a substantially more comprehensive RNA-Seq dataset was available at the time of our study (209 vs 35). We also took advantage of existing resources to further improve the annotations for these loci. These included not only their expression pattern and nucleotide conservation but also (i) the presence of potential functional elements within them (transcription factor binding and AGO binding sites), (ii) their correlation in expression with neighbouring protein coding genes, and (iii) the reported mutant phenotypes for these genes. The primary aim of these comprehensive annotations was to inform the selection of candidate lncRNA loci for follow-up experimental validation. Most importantly, we went beyond computational predictions of functionality as we assessed the in-vivo phenotypic effect of knocking-out a selection of ten intergenic lncRNAs and implemented knock-down assays to validate the observed phenotypes and putative transcript mediated function of these loci.

## lncRNAs of *C. elegans*

Our newly annotated loci bear all the hallmarks of lncRNAs in other organisms: they tend to be shorter, expressed at lower levels and have lower degree of conservation than protein coding sequences [6,7,11]. Furthermore, the GC composition of the multi-exonic lncRNAs in *C. elegans* does mirror the patterns previously observed in other animals, with increased GC content within exons relative to introns [24]. These similarities with other animal lncRNA annotations implicate *C. elegans* as a model organism that is more broadly relevant for investigation of the molecular functions of lncRNAs and the processes through which those functions are conveyed. Most importantly the wealth of resources available for *C. elegans* as a model organism, offer the opportunity to assess the *in vivo* impact of mutations within these loci.

The observation of enrichment for enhancer sequences within our lncRNA loci emphasises that the observed function of a locus could be conveyed by discrete functional DNA elements located within it, rather than by the RNA transcribed at this location. The former would imply that transcription at this location either reflects or maintains open chromatin states and that the resulting transcript would likely be biologically inconsequential, whereas the latter would imply RNA sequence-dependent functionality of the resulting transcript [10,33]. LncRNAs with transcript mediated function have been shown to act both *in cis* (Xist, [34]) and *in trans* (Paupar, [35]), whereas those whose function is transcription regulation related are expected to act in *cis* (transcriptional interference, chromatin modification at enhancers). This duality, transcription-versus transcript-mediated function, is a recurrent issue when studying lncRNAs, and only the careful experimental characterisation of each locus through knockdown, and rescue can begin to deduce the functional mechanism associated with a non-coding transcript [36].

## Phenotypic characterisation of lncRNAs

Historically, majority of *C. elegans* genes were identified through genetic screens which concurrently provide phenotypic and functional information. Mutations identified in non-coding regions of the genome as a result of genetic screens, nevertheless, have largely remained uncatalogued. With advances in genome editing methods, it is now possible to directly target non-coding regions for mutational analysis. The lncRNA annotations presented in this study, together with the detailed documentation of their expression and overlap with existing datasets, serve as a guide for the targeted analysis of these loci during animal development.

In *C. elegans,* many small non-coding RNA genes lack discernible phenotypes when deleted individually [37]. This is mostly due to redundancy between non-coding RNAs and the role such RNAs play as buffers in gene expression regulation rather than being the master regulators. The situation may be similar for the majority of lncRNAs, because their roles in the regulation of gene expression remains incompletely understood and the phenotypic characterisation of many vertebrate lncRNAs has been challenging and has provided sometimes contradictory results [38]. By using automated microscopy, we sought to capture the phenotypes associated with lncRNAs in an unbiased manner. The observed reductions in brood size and growth rate of lncRNA loci deletion mutants greatly affect the fitness of these animals, despite their otherwise normal appearance. For two of the lncRNA loci, linc-239 and linc-339, the phenotypes can be recapitulated by RNAi knock-down. We thus consider these two lncRNAs as being representative of bona fide *C. elegans* lncRNAs. However, further experiments will be required to completely rule out that these lncRNAs are not translated into functional, short polypeptides [39]. The lack of phenotypes upon RNAi knock-down of the remaining loci could be attributed to the possibility that observed phenotypes in deletion mutants arising from the removal of DNA-dependent functional elements. It is also possible that the transcripts of these loci are solely nuclear or expressed in neuronal tissues, and thus resistant to RNAi in *C. elegans* [40,41].

## Conclusions

In this study, we increased the current number of potential lncRNAs in *C. elegans* from 801 to 4001. Together with the previously identified high-confidence lncRNA loci, in total 298 loci yield evidence for possible biological functions because they display higher conservation, higher expression, higher GC content and splicing. Using genome editing and RNA interference methods, we tested the functional relevance of ten of these loci and demonstrated that six yield in vivo phenotypes when deleted. Furthermore, we showed that for at least two out of these six loci the function is likely conveyed by the RNA transcript. From our in-vivo assays, we estimate by extrapolation that 40-60% of the multi-exonic lncRNAs identified in this study might have biological roles. It will be essential to employ sensitive experimental approaches to decipher the fitness effect of such non-coding loci.

## Methods

### Intergenic lncRNA identification

A total of 209 publicly available libraries were retrieved from the SRA database (https://www.ncbi.nlm.nih.gov/sra/, Additional File 1). Reads were mapped onto the *C. elegans* (ENSEMBL release 73, WBcel235) reference genome using TOPHAT2 [42]. For each library, de novo transcripts were called using cufflinks2 [43] and the coding potential of all new intergenic transcripts was assessed using the Coding Potential Calculator (CPC, [44], score <0). All of the loci for which every transcript was deemed non-coding, were retained for further analyses as potential intergenic lncRNAs. All lncRNAs across all libraries were merged into a single annotation file using cuffcompare. We retained for final analyses only the loci longer than 200 nt, not overlapping any annotated gene and found at least 50 nucleotides away from annotated genes if located on the same strand.

### Intergenic lncRNA conservation

The nucleotide conservation of the candidate loci was assessed using the conservation tracks (PhyloP) from the UCSC database (http://hgdownload.soe.ucsc.edu/). The tracks represent the nucleotide conservation across 26 nematode species.

### Chromatin modifications, transcription binding sites and enhancers associated with lncRNAs

In order to facilitate the prioritization of lncRNAs for mutagenesis we parsed publicly available data to further improve our annotations. We intersected our annotated loci with highly occupied target regions [45], miRNA binding sites [23], transcription factor binding sites identified by modENCODE [22], and enhancers identified by Chen et al [21]. We also computed the distance to the closest protein coding gene as well as the correlation in expression between lncRNAs and their upstream and downstream flanking protein coding genes. Finally, we also reported the known phenotypes for the proteins flanking lncRNAs. Genomic locations of the respective annotations were transferred to the ce11 genome assembly using liftOver and files available on the UCSC database. Enrichment analyses were performed using the Genomic Association Test (GAT) software [46]

### Identification of transcriptional start sites

We used the 5’ end tag sequencing data from Chen et al [21] to identify the putative transcriptional start sites of the intergenic lncRNAs. We applied the same approach the authors previously applied to their data. Clusters with at least 2 reads were kept and merged if on the same strand and within 25 nucleotides of each others.

### CRISPR/Cas9 mediated deletion of lncRNA loci

lncRNA loci were deleted using either plasmid base injection [47,48] or direct protein / RNA injection methods [29], as previously described. gRNA sequences and the primers used for screening of the F2 generation animals are given in Additional file 8. Isolated deletion mutants were backcrossed once to wild type animals. For lncRNA sequences and deletions see Additional file 9; for genotyping results of deletion mutants see Additional file 10.

### Cloning of RNAi vectors

Genomic sequences corresponding to the lncRNA loci were amplified using the primers listed in Additional file 8, cloned using Gibson Assembly [49] into the L4440 vector and transformed into competent *E. coli* strain HT115 [40].

## Automated microscopy analysis

### Growth Curves

Growth curves were estimated using long-term video imaging. In short, a custom camera system was used to record backlit images of *C. elegans* from the ex-utero egg stage to the egg laying adult stage (~65 hours). To accomplish this, an imaging system was built, which allowed 12 video cameras (Flea3 3.2MP monochrome, Point Grey) to record in parallel. These were used to record images of 40 *C. elegans* nematodes in 16mm circular arenas continuously at 1Hz for ~3 days. These “mini-wells” were placed in an enclosure where temperature was maintained at 20C ±20 *mK*. The resulting movies were analyzed off-line with a custom written MATLAB script (Mathworks). Tracking was based on the Hungarian Algorithm for linear assignments, [50–52], and yielded spatial trajectories *r*(*n, t*) and time series of attributes such as the area of the 2D projection *A*(*n, t*) and the length along the centerline *l*(*n, t*), where *n* denotes the individual and *t* denotes time. Growth curve data were calculated by first taking the time average at time *s* in a window of length *w*, 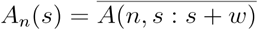. The population average in that window is then the ensemble average of the individual averages *A*(*s*) = 〈*A_n_*(*s*)〉. For these analysis wwas set at 20 minutes. Additionally, in each windows, the standard error of the mean was computed. In each set of experiments, those with mutants, and those done with RNAi bacteria, a standard growth curve was selected, the *C. elegans* N2 strain, and the *empty vector* (*ev*) bacteria respectively. For visual clarity, any other growth curves that fell within the 99% confidence interval of the “standard” curve were combined *A_J_*: 〈{*A_j_*(*s*): *p*(*H* ‡ *H*_0_(*A_j_*,*A_std_*) < 0.01}〉 Those that did not fall into this set were plotted individually (See Additional file 11 for all growth curves plotted individually with corresponding standard errors).

### Brood-size

Brood-size measurements were completed over three 24 hour intervals. First, eggs were prepared by synchronization via coordinated egg-laying. When these animals had grown to the L4 stage single animals were transferred to fresh plate (Day 0). For 3 days, each day (Days 1-3) each animal was transferred to a new plate, while the eggs were left on the old plate and allowed to hatch and grow for ~3 days, after which, the number of animals on each of these plates was counted [53] using an custom animal counting program utilizing short video recordings. Animals were agitated by tapping each plate 4 times, after this 15 frames were imaged at 1Hz and the maximum projection was used as a background image. Animals were then detected by movement using the difference image between each frame and this background image, and counted this way for 10 additional frames. The final count was returned as the mode of these counts. This system was tested on plates with fixed numbers of animals and was accurate to within 5%, comparable to human precision. Total brood size was reported then as the sum of the three days. For mutant strains, this experiment was done for 5 animals of each strain three times. For the RNAi experiment, this was done for 6 animals for each RNAi clone, also done three separate times. Data is censored for animals that crawled off of plates.

## Declarations

The authors declare that they have no competing interests.

## Funding

This work was supported by Cancer Research UK grant (C13474/A18583, C6946/A14492), the Wellcome Trust grant (104640/Z/14/Z, 092096/Z/10/Z), and The European Research Council grant (ERC, grant 260688) awarded to EAM and supported AA and EAM. DJ is supported by a Herchel Smith post-doctoral fellowship. ICN is supported by Science without Borders Full PhD scholarship (CNPq, 205589/2014-6). WH and TW are supported by a BBSRC Core Strategic Programme Grant [BB/P016774/1] and UK Medical Research Council [MR/P026028/1]. CPP is funded by the UK Medical Research Council. This research was supported in part by the NBI Computing Infrastructure for Science Group, which provides technical support and maintenance to E’s high-performance computing cluster and storage systems, which enabled us to develop this workflow.

## Authors’ contributions

AA, EAM and WH conceived the project and designed the experiments. TW and WH conducted all computational data analysis. AA and ICN generated deletion mutants. AA and DJ conducted the phenotypic analysis. DJ set up the automated microscopy system and conducted microscopy data analysis. EAM and CPP provided expertise and feedback. AA and WH wrote the manuscript with input from all authors.

## Acknowledgements

We thank the Gurdon Institute Media Kitchen for their support in providing reagents and media. We thank Lalana Songra for spending her student project time on this study.

## Additional files

Additional file 1: List of the RNA-Seq libraries used to annotate lncRNAs.

Additional file 2: Annotation of the novel lncRNAs in *C. elegans.* (GTF file)

Additional file 3: Characterization of the nucleotide sequence properties (location, composition, nucleotide conservation, functional elements) and expression of lncRNAs in *C. elegans.*

Additional file 4: Comparison of protein coding and lncRNA transcript size. (PDF file)

Additional file 5: Enrichment of lncRNAs. (PDF file)

Additional file 6: Gene structure of 10 lncRNAs and their neighbouring genes. (PDF file) Additional file 7: RT-PCR of 2 lncRNAs. (PDF file)

Additional file 8: Primers and gRNAs used in this study. (XLSX file)

Additional file 9: Sequence of 10 lncRNAs and deletions. (TXT file)

Additional file 10: Genotyping of 10 lncRNA mutants (PDF file)

Additional file 11: Extended methods for automated microscopy and phenotyping. (PDF file)

